# Transition densities and sample frequency spectra of diffusion processes with selection and variable population size

**DOI:** 10.1101/014639

**Authors:** Daniel Živković, Matthias Steinrücken, Yun S. Song, Wolfgang Stephan

**Affiliations:** Section of Evolutionary Biology, Department of Biology, Ludwig-Maximilian University Munich, Munich, Germany; Department of Statistics, University of California, Berkeley, California 94720, USA; Computer Science Division, University of California, Berkeley, California 94720, USA

## Abstract

Advances in empirical population genetics have made apparent the need for models that simultaneously account for selection and demography. To address this need, we here study the Wright-Fisher diffusion under selection and variable effective population size. In the case of genic selection and piecewise-constant effective population sizes, we obtain the transition density function by extending a recently developed method for computing an accurate spectral representation for a constant population size. Utilizing this extension, we show how to compute the sample frequency spectrum (SFS) in the presence of genic selection and an arbitrary number of instantaneous changes in the effective population size. We also develop an alternate, efficient algorithm for computing the SFS using a method of moments. We apply these methods to answer the following questions: If neutrality is incorrectly assumed when there is selection, what effects does it have on demographic parameter estimation? Can the impact of negative selection be observed in populations that undergo strong exponential growth?

## Introduction

Advances in empirical population genetics have pointed out the need for models that simultaneously account for selection and demography. Studies on samples from various species including humans (e.g., Williamson et al. 2005; Tennessen et al. 2012) and *Drosophila melanogaster* (Glinka et al. 2003; Duchen et al. 2013) have shown that demographic processes such as population size changes shape in large part the patterns of polymorphism among genomes and estimated the impact of selection on top of such underlying neutral conditions. Thus far, most theoretical papers considered selective and demographic forces independently of each other for the sake of simplicity (e.g., Stephan and Li 2007).

Theoretical studies of neutral models of time-varying population size have been accomplished within the diffusion and the coalescent frameworks. Kimura (1955a) derived the transition density function of the Wright-Fisher (WF) diffusion with a constant population size that characterizes the neutral evolution of allele frequencies over time. Shortly thereafter, Kimura (1955b) noted how to rescale time to generalize this result to a deterministically changing population size. Nei et al. (1975) derived the average heterozygosity under this general condition by applying a differential equation method, before studies on time-varying population size started to utilize the coalescent. Watterson (1984) derived the probability distribution and the moments of the total number of alleles in a sample using models of one or two sudden changes in population size. Slatkin and Hudson (1991) considered the distribution of pairwise differences in exponentially growing populations, before Griffiths and Tavaré (1994) provided the coalescent for arbitrary deterministic changes in population size. The allele frequency spectrum, which is the distribution of the number of times a mutant allele is observed in a sample of DNA sequences, has been utilized in many theoretical and empirical studies. It can be further distinguished into the allelic spectrum and the sample frequency spectrum (SFS) according to whether absolute or relative frequencies are meant. Fu (1995) derived the first- and second-order moments of the allelic spectrum for a constant population size, which has been generalized to time-varying population size by Griffiths and Tavaré (1998) and Živković and Wiehe (2008). Although deterministic fluctuations in population size are commonly considered for the interpretation of biological data, studies have also examined stochastic changes in population size (e.g., Kaj and Krone 2003).

The mathematical modeling of natural selection was mostly carried out within the diffusion framework, whereas coalescent approaches have proven analytically tedious (e.g., Krone and Neuhauser 1997). Fisher (1930) derived the equilibrium solution for the allelic spectrum of a population, which became particularly useful when Sawyer and Hartl (1992) modeled the frequencies of mutant sites via a Poisson random field approach. Kimura (1955c) employed a perturbation approach to obtain a series representation of the transition density function that is accurate for scaled selection coefficients smaller than one. However, as noted in Williamson et al. (2005), an appropriate use of this result with respect to the analysis of whole-genome data is even difficult for a constant population size. In a recent paper, Song and Steinrücken (2012) devised an efficient method to accurately compute the transition density functions of the WF diffusion with recurrent mutations and general diploid selection. This nonperturbative approach that can be applied to scaled selection coefficients substantially greater than one finds the eigenvalues and the eigenfunctions of the diffusion generator and leads to an explicit spectral representation of the transition density function. The results for this biallelic case have been extended to an arbitrary number of alleles by Steinrücken et al. (2013).

In recent years, several researchers have started to investigate the combined effect of natural selection and demography. The majority of these studies have utilized finite difference schemes to make results applicable. Williamson et al. (2005) employed such a scheme to obtain a numerical solution for the SFS for a model with genic selection and one instantaneous population size change. The authors applied this result within a likelihood-based method to infer population growth and purifying selection at non-synonymous sites across the human genome. Evans et al. (2007) investigated the forward diffusion equation with genic selection and deterministically varying population size and incorporated the effect of point mutations via a suitable boundary condition. They derived a system of ODEs for the moments of the allelic spectrum, but had to resort to a numerical scheme to make their results applicable. Gutenkunst et al. (2009) considered population substructure and selection to obtain the joint allele frequency spectrum of up to three populations by approximating the associated diffusion equation by a finite difference scheme as well. Lukić and Hey (2012) applied spectral methods that even account for a fourth population in the otherwise same setting as Gutenkunst et al. (2009). Recently, and again with respect to a single population, Zhao et al. (2013) provided a numerical method to solve the diffusion equation for random genetic drift that can incorporate the forces of mutation and selection. The authors illustrated the accuracy of their discretization approach by determining the probability of fixation in the presence of selection for both an instantaneous population size change and a linear increase in population size. In general, such methods require an appropriate discretization of grid points, which may depend strongly on the parameters. This makes it difficult, however, to predict if a particular discretization will produce accurate results.

In this study, we use the polynomial approach by Song and Steinrücken (2012) to obtain the transition density function for genic selection and instantaneous changes in population size. First, we focus on a single time period during which the population has a different size relative to a fixed reference population size. We compute the eigenvalues and the eigenfunctions of the diffusion operator with respect to the modified drift term of the underlying diffusion equation. Similarly to a constant population size, the eigenfunctions are given as a series of orthogonal functions. The eigenvalues and eigenfunctions facilitate a spectral representation of the transition density function describing the change in allele frequencies across this time period. Such transition densities for single time periods can then be folded over various instantaneous population size changes to obtain the overall transition density function for such a multi-epoch model with genic selection. After illustrating the applicability of this approach, we derive the SFS by means of the transition density function. While the transition density function proves useful for the analysis of time-series data that are mostly gathered from species with short generation times as bacteria (e.g., Lenski 2011) but also from species with long generation times (Steinrücken et al. 2014), the SFS can also be applied to whole-genome data collected at a single time point. As an alternative approach to employing the transition density function for the SFS, we modify the method of moments by Evans et al. (2007) to efficiently compute allele frequency spectra for genic selection, point mutations and piecewise changes in population size.

We then employ a maximum likelihood method to estimate the demographic and selective parameters of a given bottleneck model. After examining the accuracy of parameter estimation, we discuss how the estimates change when selection is ignored or a simpler demographic model is assumed. In this context, we investigate the demography of an African population of *Drosophila melanogaster* (Duchen et al. 2013) by assuming both a selective and a neutral model. Furthermore, we answer an other, important question arising in human population genetics (Tennessen et al. 2012): Can the impact of negative selection be observed in populations that undergo strong exponential growth? We investigate, how strong selection would have to be to leave a signature in the SFS.

## The transition density function for genic selection and piecewise-constant population sizes with *K* epochs

### Model and notation

We assume that the diploid effective population size changes deterministically, with *N*(*t*) denoting the size at time *t*. Here, time is measured in units of 2*N*_ref_ generations, where *N*_ref_ is a fixed reference population size. Unless stated otherwise, the initial population size will be used as the reference population size in the various numerical examples. In the diffusion limit, the relative population size *N*(*t*)/*N*_ref_ converges to a scaling function which we denote by *ρ*(*t*).

We assume the infinitely-many-sites model (Kimura 1969) with *A*_0_ and *A*_1_ denoting the ancestral and derived allelic types, respectively. The relative fitnesses of *A*_1_/*A*_1_ and *A*_1_/*A*_0_ genotypes over the *A*_0_/*A*_0_ genotype are respectively given by 1 + 2*s* and 1 + *s*. The population-scaled selection coefficient is denoted by *σ* = 2*N*_ref_ · *s*. The frequency of the derived allele *A*_1_ at time *t* is denoted by *X*_*t*_. Let ƒ be a twice continuously differentiable, bounded function over [0,1]. The backward generator of a time-inhomogeneous one-dimensional WF diffusion process on [0,1] is denoted by *ℒ*, which acts on f as

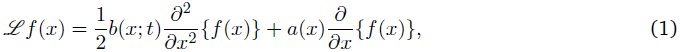

where the diffusion and drift terms are given by *b*(*x*;*t*) *= x*(1 − *x*)/*ρ*(*t*) and *a*(*x*) = *σx*(1 − *x*), respectively. The dependence of the diffusion term on time introduces considerable challenges to obtaining analytic results. To gain insights, we here focus on the case where *ρ* is piecewise constant. In this case, the diffusion and drift terms differ by a constant factor within each piece, thus simplifying the analysis.

Throughout, we assume that *ρ* has *K* constant pieces (or epochs) in the time interval [*τ*_0_,*τ*). The change points are denoted by *t*_1_,…, *t*_*K*−1_, and for convenience we define *t_0_* = *τ_0_* and *t_K_* = *τ*. Then, for *t*_*i*_ ≤ *t* < *t*_*i*+1_, with 0 ≤ *i* ≤ *K* − 1, we assume *ρ*(*t*) = *c*_*i*_, where *c*_*i*_ is some positive constant. For the epoch *t_i_* ≤ *t* < *t*_*i*+1_, the diffusion term is thus given by *b*(*x*) = *x*(1 − *x*)/*c_i_* and the corresponding generator is denoted by ℒ^*i*^. The scale density *ξ_i_* (Karlin and Taylor 1981, Ch. 15) for the epoch is given by

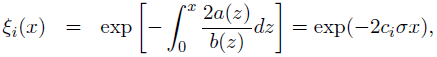

while the speed density *π_i_* is given (up to a constant) by

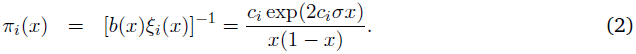

Given real-valued functions ƒ and *g* on [0,1] that satisfy appropriate boundary conditions and are square integrable with respect to some real positive density *h*, we use 〈ƒ,*g*〉_*h*_ to denote

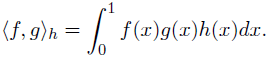

### **The transition density within each epoch** [*t*_*i*_,*t*_*i*+1_)

For the epoch [*t*_*i*_,*t*_*i*+1_), let the transition density be denoted by *p*_*i*_(*t*;*x*,*y*), where *t* ∈ [*t*_*i*_,*t*_*i*+1_), *X*_*t*_*i*__ = *x* and *X*_*t*_ = *y*. Under the initial condition *p*_*i*_(*t*_*i*_; *x*, *y*) = *δ*(*x* − *y*), the spectral representation of *p*_*i*_(*t*; *x*, *y*) is given by

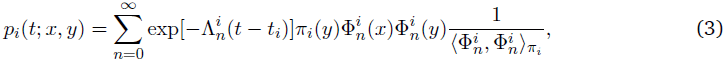

where 
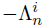 and 
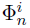 are the eigenvalues and eigenfunctions of **ℒ*^i^*, respectively. That is,

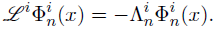

It can be shown that the eigenvalues are all real and non-positive. Furthermore,

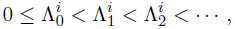

with 
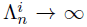 as *n* → ∞. The associated eigenfunctions 
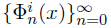 form an orthogonal basis of *L*^2^([0,1],*π_i_*), the space of real-valued functions on [0,1] that are square integrable with respect to the speed density *π_i_*, defined in (2).

Song and Steinrücken (2012) recently developed a method for finding 
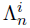 and 
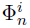 in the case of *c_i_* = 1. We will give a brief description of their method and modify it accordingly to incorporate an arbitrary *c_i_* > 0. Let 
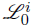 denote the diffusion generator under neutrality (i.e., *σ* = 0). The eigenfunctions of 
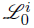 are modified Gegenbauer polynomials 
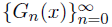 (cf. *Appendix*), and the corresponding eigenvalues are 
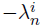, with

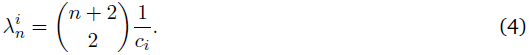

Similar to Song and Steinrücken (2012), define 
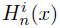 as

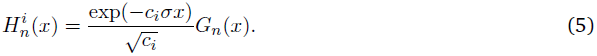

Then, 
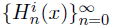 form an orthogonal system with respect to the weight function *π_i_*(*x*). By directly applying the full generator *ℒ^i^* to 
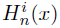, we observe that 
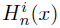 are not eigenfunctions of *ℒ^i^*. Instead, we obtain

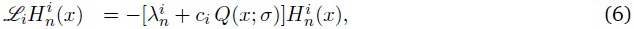

where *Q*(*x*; *σ*) = 1/2 · *σ*^2^(*x*1 − *x*). However, since both 
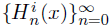 and 
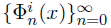 are orthogonal with respect to the same weight function *π_i_(*x*)*, and 
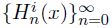 form a basis of *L^2^*([0,1],*π_i_*), we can represent 
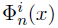 as a linear combination of 
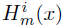:

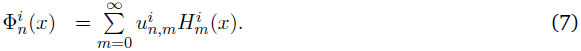

Furthermore, the fact that 
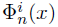 is an eigenfunction of *ℒ^i^* with eigenvalue 
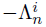 implies that 
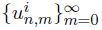 and 
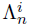 satisfy the following equation:

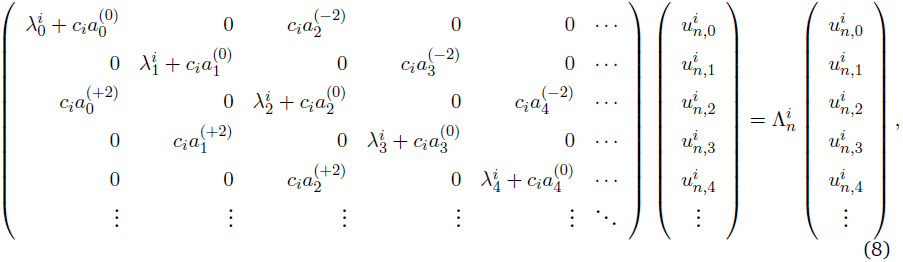

where 
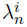 is as defined in (4) and 
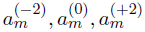 are known constants that depend on *σ* and *m* (cf. Song and Steinrücken 2012 for details).

The transition density expansion (3) can be obtained by numerically solving the eigensystem (8). Denote the infinite-dimensional matrix on the left hand side of (8) by *W_i_*. The eigenvalues 
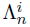 of *W_i_* correspond (up to a sign) to the eigenvalues of *ℒ^i^*, and the associated eigenvectors 
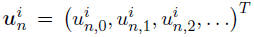 of *W_i_* determine the eigenfunctions of *ℒ^i^* via (7). Let 
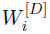 denote the *D* × *D* matrix obtained by taking the first *D* rows and *D* columns of *W_i_*, and let 
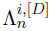 and 
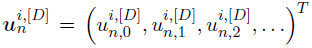 denote the eigenvalues and eigenvectors of 
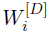, respectively. The truncated eigensystem

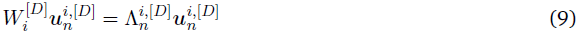

can then be used to approximate (8). This finite-dimensional linear system can be easily solved numerically. Since the truncated versions of the eigenvalues and eigenvectors converge rapidly as *D* increases, an accurate approximation of the transition density (3) can be efficiently obtained. The rapid convergence behavior of the eigenvalues is illustrated in Figure 1. As one would expect, the truncation level *D* required for convergence is higher when modeling a large population (cf. Figure 1b) compared to the basic selection model (cf. Figure 1a), while convergence is fast in a model with smaller population size (cf. Figure 1c). This is because the necessary truncation level correlates with the effective strength of selection, which is higher in large populations and lower in small populations. Therefore, for a fixed selection coefficient *s*, large populations are computationally more demanding than small populations.

**Figure 1.**
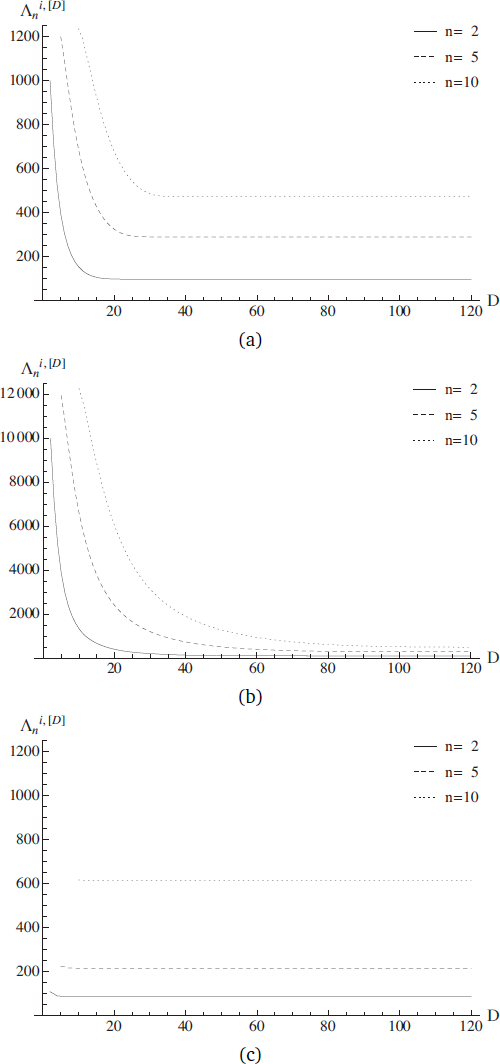
Convergence of the eigenvalues 
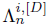 with increasing truncation level *D* for (a) a constant population size (c_*i*_ = 1), (b) a large population (c_*i*_ = 10) and (c) a small population (c_*i*_ = 1/10). The eigenvalues are plotted for three values of *n* and a scaled selection coefficient of *σ* = 100 in each panel.

### **The transition density for the entire period** [*τ*_0_, *τ*) **with** *K* **epochs**

Suppose *X_τ_*_0_ *= x* and *X_τ_* = *y*. The transition density *p*(*τ*_0_,*τ*;*x*,*y*) for the entire period [*τ*_0_,*τ*) is obtained by combining the transition densities for the *K* epochs as follows:

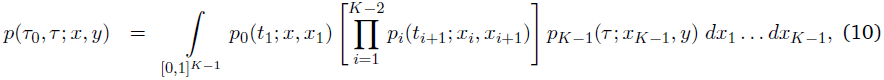

where *x_i_* denotes the allele frequency at the change point *t_i_*. Using (3), we can write (10) as

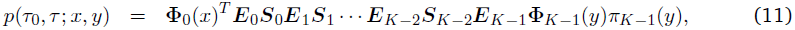

where 
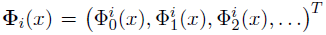 is an infinite-dimensional column vector, while *E_i_* and *S_i_* are infinite-dimensional matrices defined as

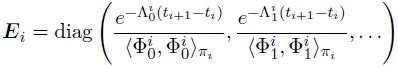

and

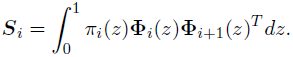

In general, *S_i_* is not a diagoNal Matrix Since 
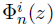 and 
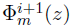 are not orthogonal with respect to
π_*i*_(*z*) if *c*_*i*_ = *c*_*i*+1_. In the *Appendix*, we show that the entry (*n, m*) of *S*_*i*_ is given by

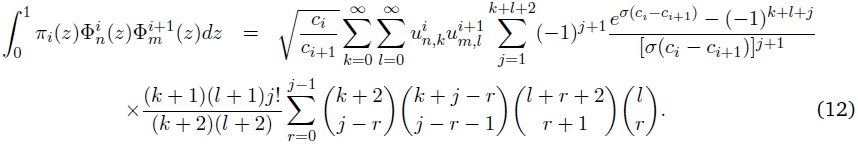

Note that the last line of (12) does not depend on *n* or *m*, so it needs to be computed only once. The overall computational time for evaluating *p*(*τ*_0_,*τ*;*x*,*y*) scales linearly with the number *K* of epochs.

To better understand the joint impact of selection and demography on the transition density function, we consider two scenarios, where *p*(0,*τ*;*x*,*y*) is simply denoted as *p*(τ;*x*,*y*). Figure 2 illustrates the density in a scenario in which the selection coefficient is fixed and various *K*-epoch demographic models are considered. In comparison to the case of a constant population size (cf. Figure 2a), an instantaneous expansion (cf. Figure 2b) narrows the distribution around the mean, whereas an additional phase of a reduced population size (cf. Figure 2c) increases the variance relative to a population of a constant size. Figure 3 illustrates the same scenarios with a fixed transition time and varying selection coefficients.

**Figure 2.**
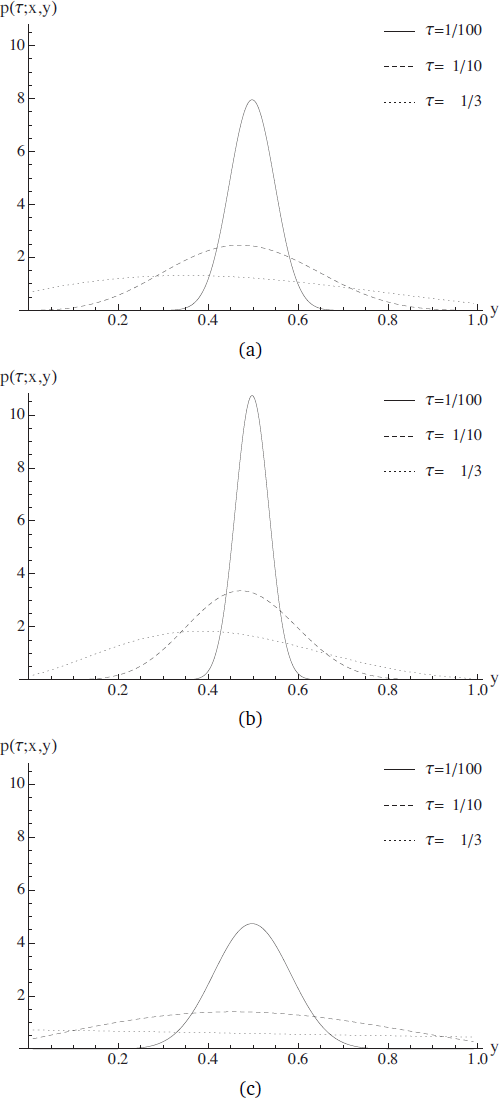
Transition densities for various transition times *τ* and a fixed selection coefficient *σ* = − 1. In all cases, we set *x* = 1/2 and *D* = 100. (a) A single-epoch model (*K* = 1), a constant population size with *c*_0_ = 1 (b) A two-epoch model (*K* = 2), with an instantaneous expansion (*c*_0_ = l,*c*_1_ = 10, *t*_1_ = τ/2). (c) A three-epoch model (*K* = 3), with a population bottleneck followed by an expansion (*c*_0_ = l,*c*_1_ = 1/10, *c*_2_ = 10,*t*_1_=r/4,*t*_2_=τ/2).

**Figure 3.**
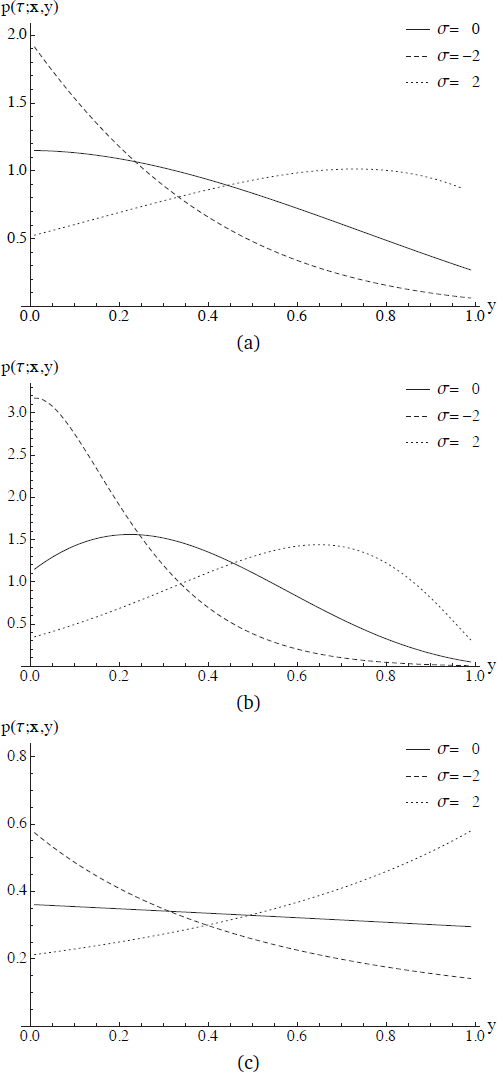
Transition densities for various selection coefficients *σ* and a fixed transition time *τ* = 1/2. In all cases, we set *x* = 1/3 and *D* = 100. (a) A single-epoch model (*K* = 1), a constant population size with *c*_0_ = 1. (b) A two-epoch model (*K* = 2), with an instantaneous expansion (*c*_0_ = 1, *c*_1_ = 10, *t*_1_ = τ/2). (c) A three-epoch model (*K* = 3), with a population bottleneck followed by an expansion (*c*_0_ = 1, *c*_1_ = 1/10, *c*_2_ = 10,*t*_1_=τ/4,*t*_2_=τ/2).

## The sample frequency spectrum (SFS)

### The transition density function approach

The transition density function derived in the previous section can be employed to obtain the SFS of a sample. Consider a sample of size *n* obtained at time *t* = *τ*. The probability that the *A*_1_ allele with frequency *x* at time *t* = *τ*_0_ is observed *b* times in the sample is (Griffiths 2003)

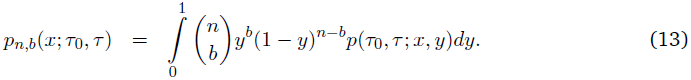

For piecewise-constant population size models with *K* epochs, a spectral representation of *p*(τ_0_,*τ*;*x*,*y*) can be found via (11) and evaluating (13) involves computing the integral 
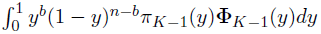. For *l* ≥ 0, using (2), (5), and (7), we obtain

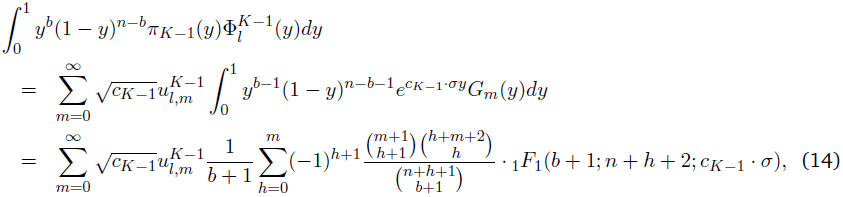

where 
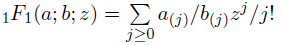 is the confluent hypergeometric function of the first kind. The descending factorials *d*_(*j*)_ are defined in the *Appendix*.

The sample frequency spectrum (SFS) *q_n_*,*_b_*(τ) is the probability distribution on the number *b* of mutant alleles in a sample of size *n* taken at time *τ*, conditioned on segregation. For 1 ≤ *b* ≤ *n* − 1, *q_n_*,*_b_*(τ) is given by

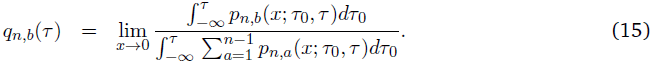

In (15), the SFS at a single site is obtained by averaging over sample paths. This is equivalent to the frequency spectrum distribution over a large number of independent mutant sites in the Poisson random field model of Sawyer and Hartl (1992). Using (11), (12), (13), and (14), we can approximate (15) numerically. If it is unknown which allele is derived, a folded version of (15) can be obtained as [*q*_*n*,*b*_ + *q*_*n*,*n*−*b*_]/(1 + δ_*b*,*n*−*b*_), where δ_*b*,*n*−*b*_ denotes the Kronecker delta.

### A method of moment approach

As detailed above, the transition density function can be employed to obtain the SFS. However, the specific solution for the transition density is not required to obtain the less complex and thus computationally less demanding SFS. Here, we utilize the work of Evans et al. (2007) to develop an efficient algorithm for computing the allele frequency spectrum in the case of genic selection and piecewise-constant population sizes.

Suppose mutations arise at rate *θ*/2 (per sequence per 2*N*_ref_ generations) and according to the infinitely-many-sites model (Kimura 1969). Evans et al. (2007) use the forward diffusion equation to describe population allele frequency changes and introduce mutations by an appropriate boundary condition. Slightly modifying their notation, we use ƒ(*y*, *t*)*dy* to denote the expected number of sites where the mutant allele has a frequency in (*y*, *y* + *dy*), with 0 < *y* < 1, at time *t*. The forward equation is

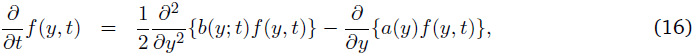

where the diffusion term *b*(*y*; *t*) = *y*(1 − *y*)/*ρ*(*t*), the drift term *a*(*y*) = *σy*(1−*y*), the scaled selection coefficient *σ*, and the population size function *ρ*(*t*) are defined as before. The influx of mutations is incorporated into this process via the boundary conditions

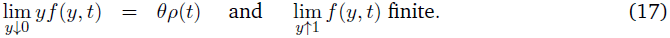

The resulting polymorphic sites follow the dynamics of (16) thereafter. Note that this differs from the diffusion process studied in the previous section, as the influx of mutations is now explicitly modeled.

Again, it is analytically more practical to consider the corresponding backward equation, which is obtained by setting *g*(*y*,*t*):= *y*(1 − *y*)ƒ(*y*, *t*). This substitution transforms the forward equation for ƒ(*y*, *t*) into a backward equation for *g*(*y*, *t*), which is essentially given by (1) up to the sign of the drift term. Evans et al. (2007) derived a coupled system of ordinary differential equations (ODEs) for the moments 
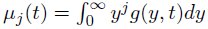

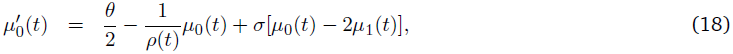

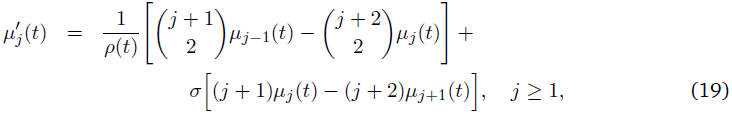

where 
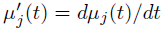. A similar system of ODEs was derived and solved by Kimura (1955a) for a neutral scenario with a constant population size and without mutations. For σ = 0, the above system is finite and can be solved explicitly (Živković and Stephan 2011). In the case of selection (σ ≠ 0), on the other hand, the system is infinite and obtaining an explicit solution for an arbitrary *ρ* is a challenging problem, even if the system is truncated by setting *µ_j_*(*t*) = 0 for *j* ≥ *D*.

From now on, assume *µ_j_* (*t*) ≡ 0 for *j* ≥ *D* and rewrite the truncated system of ODEs in matrix form as

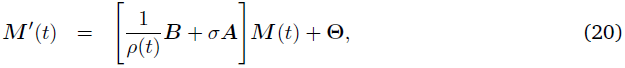

where 
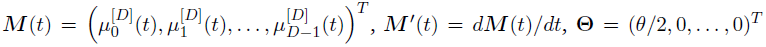 are *D*-dimensional column vectors, and ***B*** = (*b_kl_*) and ***A*** = (*a_kl_*) are *D* × *D* matrices with entries

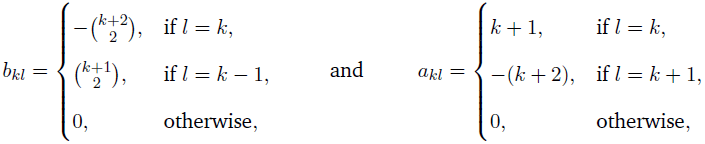

for 0 ≤ *k*, *l* ≤ *D* − 1. The formal solution of (20) cannot be written in terms of a matrix exponential but only as a Peano-Baker series (Baake and Schlagel 2011) for arbitrary *ρ*, which can be numerically quite demanding. Therefore, we focus on the case of piecewise constant *ρ* and develop an efficient method to solve the truncated system of ODEs.

We first consider *ρ*(*t*) ≡ *c*_0_ (i.e., a constant population size), for which the solution of (20) takes the form of a matrix exponential given by

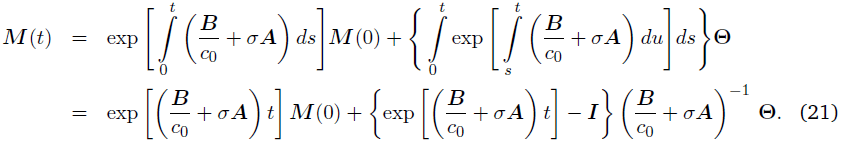

Let –λ_*k*_, (*l*_k,0_, …, *l*_*k*,*D*‒1_), and (*r*_0,*k*_, …, *r*_*D*‒1,*k*_)^*T*^ respectively denote the eigenvalues, row eigenvectors, and column eigenvectors of *B*/*c*_0_ + *σ* ***A***. Then, (21) implies

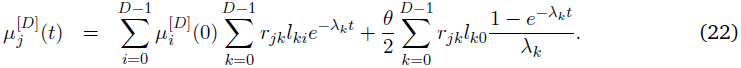

It is intractable to find closed-form expressions of −*λ_k_*, l_*ki*_, and *r_jk_*, but, for a given truncation level *D*, they can be computed numerically. Depending on the details of the model under consideration, it might be more efficient to solve (21) numerically rather than applying the more analytic form given in (22).

We now investigate the equilibrium solution of (22), since it can be applied as an initial condition in a model in which the population size remains constant over a longer period of time before instantaneous population size changes occur. Assuming that all alleles are monomorphic at time zero, i.e. 
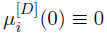, and letting *t* → ∞, we obtain the moments at equilibrium as

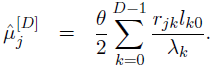

For *D* sufficiently large, this result is numerically close to the exact solution µ^*_j_*. The latter can also be obtained as follows. The equilibrium population frequency spectrum is given by (Fisher 1930)

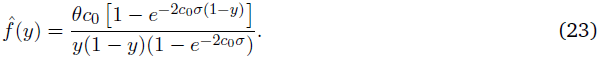

The sampled version can be easily found via binomial sampling as in (13):

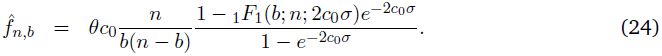

For *σ* ≠ 0, the moments µ^*_j_*of *ĝ*(*y*) = *y*(1 − *y*)ƒ^(*y*) are given by

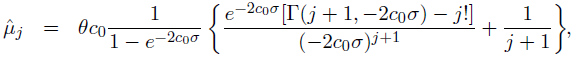

where 
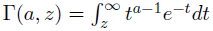 is the incomplete gamma function.

Now, consider the piecewise-constant model with *K* epochs in the time interval [*τ*_0_,τ] defined earlier. For *t_i_* ≤ *t* <*t*_*i*+1_,

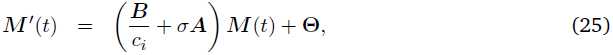

which can be solved as in (21). For *τ* > *t*_*K*−1_,

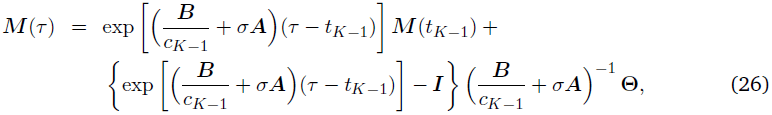

where *M*(*t_i_*), for 1 ≤ *i* ≤ *K* − 1, is recursively given by

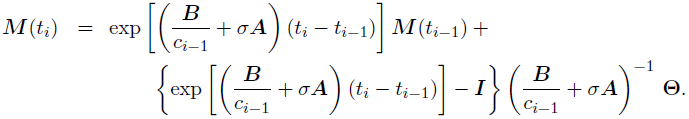

The initial condition *M*(*t*_0_) is either chosen as the equilibrium solution described above or the zero vector, which corresponds to the case of all loci being monomorphic at time *t*_0_ = *τ*_0_.

The accuracy of the above framework depends on how fast the truncated moments 
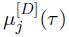 converge to zero as *D* increases. In general, the truncated moments converge faster for negative than for positive *σ*, and for instantaneous declines compared to instantaneous expansions (cf. Figure 4). For a large positive *σ*, a higher truncation level *D* may be required to achieve the desired accuracy. Finally, the allelic spectrum *f_n_*_,*b*_(*τ*), for 1 ≤ *b* ≤ *n* − 1, of a sample of size *n* taken at time *τ* can be obtained from the moments *µ_j_* (*τ*) via (26) and by using the relationship

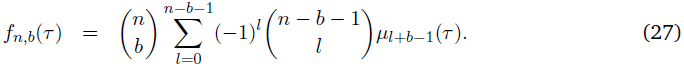

**Figure 4.**
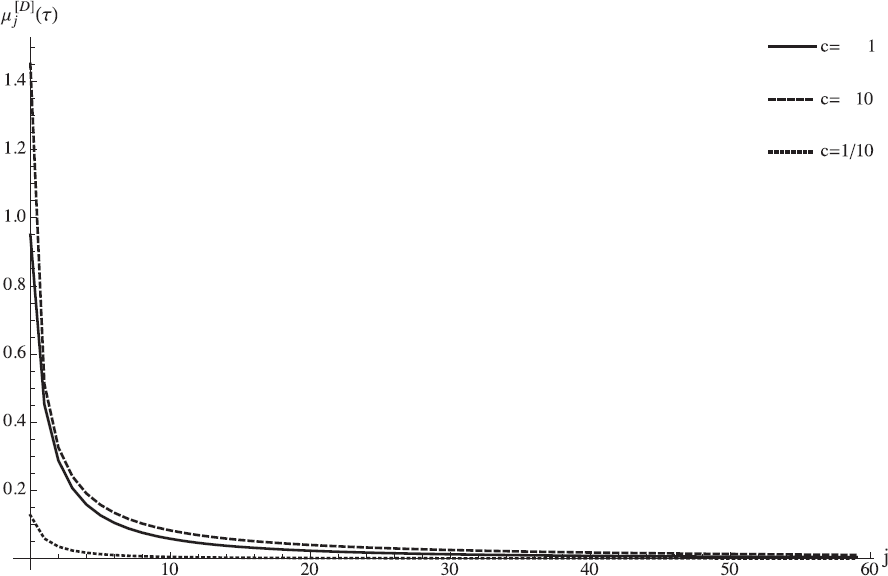
Convergence of the moments 
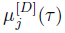 as *j* increases, with *D* = 500, τ = 1/4 and *σ* = 10. The moments are in equilibrium until time zero, when the population size is either kept constant to *c* = 1 or instantaneously changed to *c* = 10 or *c* = 1/10.

The SFS *q_n,b_*(*τ*) at time *τ* is then given by

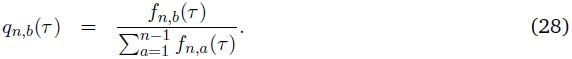

The joint impact of a population bottleneck and selection on the SFS is illustrated in Figure 5 for various points in time. As expected, negative and positive selection result in a skew of the SFS towards low- and high-frequency derived variants, respectively, when compared to a model without selection, across all sampling times. Moreover, this skew varies in intensity at different points in time. In the neutral demographic model (cf. Figure 5b), the relative frequency of singletons at time *τ*_3_ is higher than at time *τ*_4_, whereas under the same demographic model with negative selection (cf. Figure 5c) this relation is inverted. This is because the amount of singletons that is caused by demographic forces decreases after the expansion from *τ*_3_ to *τ*_4_, while negative selection is still increasing the low-frequency derived classes in this time interval.

**Figure 5.**
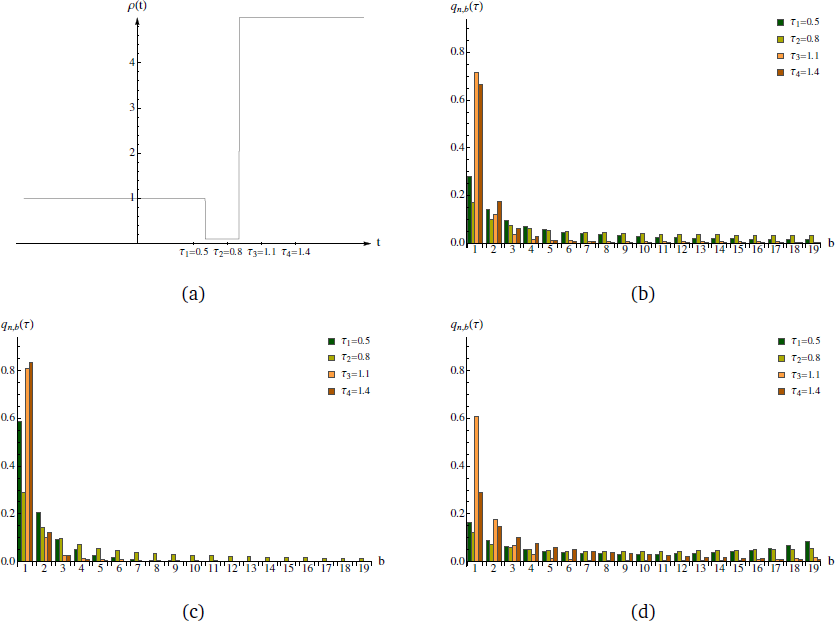
(a) The relative population size, *ρ*(*t*), is initially 1 and changes instantaneously to 1/10 and 5 at times 6/10 and 9/10, respectively. The SFS of a sample of size 20 are plotted for this demography (b) without selection, (c) negative selection of *σ* = −2 and (d) positive selection of *σ* = 10. The times of sampling are illustrated in (a) and the bars are accordingly displayed from the left to the right.

## Applications

Here, we discuss some biologically relevant questions that can be addressed using our theoretical framework. This section consists of the following three parts:

1. We first consider models with negative selection and bottlenecks of medium strength at different time points. We examine the SFS under such models and try to estimate the demographic parameters while taking selection into account. We also carry out demographic inference while ignoring selection. Whereas the former demonstrates how well the demographic and selective parameters can be estimated jointly, the latter mimics the common practice of assuming genome-wide polymorphic sites as putatively neutral (due to the difficulty of jointly estimating the impact of selection and demography using existing tools). We finally examine the consequences of assuming a too simple underlying demography on parameter estimation.

2. We then analyze an African sample of *Drosophila melanogaster* to investigate its demographic history and possible selective effects.

3. Lastly, we examine a model of strong exponential population growth (mimicking human evolution) and superimpose negative selection of various strengths to understand if and when selection can be inferred for such a model.

Throughout, the first population size change will occur after the allele frequencies have reached an equilibrium according to (24).

### Joint inference of population bottleneck and purifying selection

#### A maximum likelihood approach

Under the assumption that the considered sites are independent, the log-likelihood of a model *M* given data *D* is 
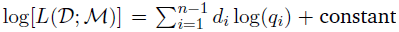, where *d_i_* is the observed number of sites at which the derived allele occurs *i* times in the sample, and *q_i_* is the probability that the derived allele occurs *i* times in the sample at a segregating site under model *M* (e.g., Wooding and Rogers 2002). Recall that *q_i_* can be either obtained via the transition density function or the method of moment approach.

Consider the bottleneck model illustrated in Figure 6. Note that the present relative size *c_S_* is fixed to 1, i.e., here the present population size is used as the reference population size *N*_ref_. First, we consider the scenario where the ancestral population size *c*_0_ prior to the bottleneck is allowed to vary. In this case, the model has five free parameters: c_0_, the initial population size; *c_B_*, the population size during the bottleneck; *t_B_*, the duration of the bottleneck; *t_S_* = *τ* − *t_B_*, the time since recovery from the bottleneck; and *σ*, the scaled selection coefficient. We then also consider the scenario where the ancestral population size is the same as the present population size, i.e., *c*_0_ = *c_S_*, resulting in a model with four free parameters.

**Figure 6.**
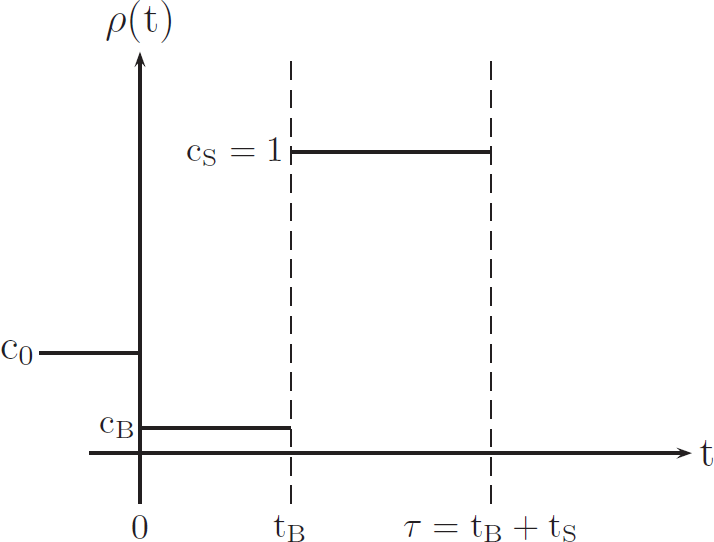
The population is constant in size before being instantaneously changed to relative size *c_B_* at time zero. Then, another jump to relative population size *c_s_* follows at time *t_B_*, before a sample is taken at time *τ* = *t_B_* +*t_S_*.

We adopted a grid search in our estimation procedure, with *σ* ∈ [−10,0] and *c_B_*,*t_B_*,*t_S_* ∈ [0.001,1]. For the 5-parameter model, *c*_0_ was chosen from the range [0.01,10]. In total, 110,000 grid points were chosen in the selected case and 10,000 in the neutral case. Note that the grid search also accounts for models of one or two successive instantaneous population expansions. For the 4-parameter model, 11,000 grid points were chosen in the selected case and 1000 in the neutral case. The grid points are summarized in Table 1.

**Table 1.** Grid values chosen for each parameter in our optimization procedure The underlying bottleneck model is illustrated in Figure 6. Grid values c_0_ ≥ *c_B_* are considered for the 5-parameter model, whereas *c*_0_ = *c_s_* in the 4-parameter model. The grid values for the remaining parameters are applied in both scenarios. The ratio of two consecutive values remains constant between (and including the) two subsequent bold entries.

|  |  |  |  |  |  |  |  |  |  |  |  |
| --- | --- | --- | --- | --- | --- | --- | --- | --- | --- | --- | --- |
| $c_0$ | **0.011** | 0.023 | **0.05** | **0.1** | 0.224 | **0.5** | **1** | 2.154 | 4.642 | **10** | |
| $\sigma$ | **-10** | -5.848 | -3.420 | **-2** | -1.260 | -0.79 | **-0.5** | -0.292 | -0.171 | **-0.1** | **0** |
| $c_B$ | **0.001** | 0.0022 | **0.005** | 0.011 | 0.023 | **0.05** | **0.1** | 0.224 | **0.5** | **1** | |
| $t_B$ | **0.001** | 0.0022 | **0.005** | 0.011 | 0.023 | **0.05** | **0.1** | 0.224 | **0.5** | **1** | |
| $t_S$ | **0.001** | 0.0022 | **0.005** | 0.011 | 0.023 | **0.05** | **0.1** | 0.224 | **0.5** | **1** | |

#### Estimation of bottleneck and selection parameters

We first evaluated the SFS for a sample of size *n* = 50 in the following twelve scenarios, all with *c_S_* = 1 and *σ* ∈ {0, −1/2, −2}:

1. Constant population size (i.e., *c*_0_ = *c_B_* = *c_S_* = 1).

2. Bottleneck models with c_0_ = 1/2, *c_B_*= 1/10, *t_B_* = 1/10, and *t_S_* ∈ {1/200,1/20,1/2}.

First, to test how well the demographic and selective parameters can be estimated jointly from sampled data, we focused on the bottleneck demography with *t_S_* = 1/20 and considered two scenarios: The neutral case (*σ* = 0) and the selected case with *σ* = −2. To mimic the limited availability of independent polymorphic sites across the genome, we sampled 10,000 sites according to the SFS for the two chosen scenarios, and repeated this procedure 200 times. For each of these 200 datasets, we maximized the log-likelihood over the grid of parameter values described earlier, assuming (A1) neutrality when the true model has *σ* = 0, (A2) neutrality when the true model has *σ* = −2, (A3) presence of selection when the true model has *σ* = −2, and (A4) presence of selection when the true model has *σ* = 0.

The estimated parameters are shown in Table 2. For inference under correct model assumptions (A1 and A3), the median estimates are equal to the true parameters. When selection is ignored although present in the dataset (A2), the ancestral population size (ĉ_0_) and the duration of the bottleneck (*t*^*_B_*) are underestimated, whereas the bottleneck size (*ĉ_B_*) and the time since the bottleneck (*t^_S_*) are accurately estimated. When the true model is neutral but the inference procedure allows for selection (A4), a neutral demographic model is accurately inferred. We calculated likelihood-ratio statistics for each of the 200 datasets to compare the two nested models of selection and neutrality. The null hypothesis of neutrality can be rejected at the 5% significance level with a power of 55%.

**Table 2.** Parameter estimation results based on 10,000 sampled sites SFS were computed for the true parameters and the demography illustrated in Figure 6 (*c*_0_ = 1/2, *c_s_* = 1). Then, 10,000 sites were sampled according to the SFS of the neutral and the selective scenario, and this procedure was repeated 200 times each. The log-likelihood values were maximized over the parameter spaces as specified in the main text, and the table reports the median, the 0.05 and the 0.95 quantiles. The four cases correspond to assuming (Al) neutrality when *σ* = 0, (A2) neutrality when *σ* = −2, (A3) presence of selection when *σ* = -2, and (A4) presence of selection when *σ* = 0.

| | | $\hat{c}_0$ | $\hat{\sigma}$ | $\hat{c}_B$ | $\hat{t}_B$ | $\hat{t}_S$ |
| --- | --- | --- | --- | --- | --- | --- |
| True parameters | | 0.5 | 0 or $-2$ | 0.1 | 0.1 | 0.05 |
| (A1) | 5% | 0.5 |  | 0.1 | 0.1 | 0.05 |
|  | Median | 0.5 |  | 0.1 | 0.1 | 0.05 |
|  | 95% | 0.5 |  | 0.1 | 0.1 | 0.05 |
| (A2) | 5% | 0.22 |  | 0.02 | 0.005 | 0.05 |
|  | Median | 0.22 |  | 0.1 | 0.05 | 0.05 |
|  | 95% | 0.22 |  | 0.1 | 0.05 | 0.05 |
| (A3) | 5% | 0.22 | $-2$ | 0.05 | 0.01 | 0.05 |
| | Median | 0.5 | $-2$ | 0.1 | 0.1 | 0.05 |
|  | 95% | 0.5 | 0 | 0.1 | 0.1 | 0.05 |
| (A4) | 5% | 0.5 | $-0.5$ | 0.1 | 0.001 | 0.05 |
|  | Median | 0.5 | 0 | 0.1 | 0.1 | 0.05 |
|  | 95% | 2.15 | 0 | 0.1 | 0.1 | 0.05 |

We further analyzed all twelve scenarios using the expected SFS directly, assuming that the amount of data is sufficiently large such that the observed SFS has converged to the expected value. Our goal in this case is to study the effect of model misspecification on parameter estimation; specifically, assuming selection when the true model is neutral or assuming neutrality when there is selection. In the former case, the maximum likelihood estimates always coincided with the true parameters. Therefore, it is useful to allow for selection in an analysis even when putatively neutral regions are considered. In the latter case, our results are summarized in Table 3. For a constant population size, two rather old instantaneous expansions are estimated. For the bottleneck models, ignoring selection leads to the largest errors for the most recent bottleneck and *σ* = −1/2 and the least recent bottleneck and *σ* = −2, for which an instantaneous expansion is estimated. Interestingly, the time since the bottleneck was robustly estimated in many cases.

**Table 3.** Parameter estimation results based on the expected SFS assuming neutrality when the true model is under selection SFS were computed for the following demographic scenarios and selection coefficients. In terms of the demography, either a constant population size was assumed, or a bottleneck model according to Figure 6 with parameters *c*_0_ = 1/2, *c_B_* = 1/10, *c_s_* = 1, *t_B_* = 1/10 and *t_s_* = 1/200, 1/20 or 1/2. The selection coefficients are *σ* = −1/2 and −2. The parameter estimates were obtained according to the procedure and the parameter spaces described in the main text and by assuming neutrality in each case. In the first row, and in the forth row, second column, we obtained *ĉ_B_* = 1, i.e. an instantaneous expansion occurs as the only size change *t^^^_B_* + *t^^^_s_*before sampling.

| Selection coefficient | $\sigma = -1/2$ | $\sigma = -2$ |
| --- | --- | --- |
| Demographic model | $(\hat{c}_0, \hat{c}_B, \hat{t}_B, \hat{t}_S)$ | $(\hat{c}_0, \hat{c}_B, \hat{t}_B, \hat{t}_S)$ |
| Constant population size | $(0.500, 1.00, 1.10 - \hat{t}_S, \hat{t}_S)$ | $(0.100, 1.000, 0.523 - \hat{t}_S, \hat{t}_S)$ |
| Bottleneck with $t_S = 1/200$ | $(0.224, 0.05, 0.05, 0.002)$ | $(0.224, 0.100, 0.050, 0.005)$ |
| Bottleneck with $t_S = 1/20$ | $(0.500, 0.10, 0.10, 0.050)$ | $(0.224, 0.100, 0.050, 0.050)$ |
| Bottleneck with $t_S = 1/2$ | $(1.000, 0.05, 0.10, 0.500)$ | $(0.100, 1.000, 0.324 - \hat{t}_S, \hat{t}_S)$ |

To assess the impact of assuming a slightly simplified model for parameter estimation, we carried out an analogous study where the ancestral population size *c*_0_ was incorrectly assumed to equal the current size *c_s_* = 1, while the true model had *c*_0_ = 1/2 and *c_s_* = 1. For the resampling analysis, we considered the same bottleneck scenarios as before with *σ* = 0 or −2, and maximized the log-likelihood values over a grid in the parameter space (as described earlier) for each of the 200 simulated datasets each containing 10,000 polymorphic sites. The parameter estimates are shown in Table 4. The time since the bottleneck (t^*s*) is accurately estimated irrespective of correct or wrong assumptions regarding selection. Incorrectly assuming *c*_0_ = *c_s_* results in either an over-estimation of the duration of the bottleneck (t^_*B*_) in most of the cases (A1−A3) or an inference of selection when *σ* = 0 (A4). Selection was poorly estimated even under (A3).

**Table 4.** Parameter estimation results based on 10,000 sampled sites when the ancestral population size c_0_ is incorrectly assumed to equal the current size *c_s_*, while the true model has *c*_0_ = 1/2 and *c_s_* = 1. SFS were computed for the true parameters and the demography illustrated in Figure 6 (*c*_0_ = 1/2, *c_s_* = 1). Then, 10,000 sites were sampled according to the SFS of the neutral and the selective scenario, and this procedure was repeated 200 times each. The log-likelihood values were maximized over the 4-parameter space (where *c*_0_ = *c_s_* is assumed), and the table reports the median, the 0.05 and the 0.95 quantiles. The four cases correspond to assuming (Al) neutrality when *σ* = 0, (A2) neutrality when *σ* = −2, (A3) presence of selection when *σ* = −2, and (A4) presence of selection when *σ* = 0.

| | | $c_0$ | $\hat{\sigma}$ | $\hat{c}_B$ | $\hat{t}_B$ | $\hat{t}_S$ |
| --- | --- | --- | --- | --- | --- | --- |
| True parameters | | 0.5 | 0 or $-2$ | 0.1 | 0.1 | 0.05 |
| (A1) | 5% |  |  | 0.1 | 0.22 | 0.02 |
|  | Median |  |  | 0.1 | 0.22 | 0.05 |
|  | 95% |  |  | 0.22 | 0.5 | 0.05 |
| (A2) | 5% |  |  | 0.1 | 0.22 | 0.05 |
|  | Median |  |  | 0.1 | 0.22 | 0.05 |
|  | 95% |  |  | 0.22 | 1 | 0.05 |
| (A3) | 5% | | $-0.79$ | 0.1 | 0.22 | 0.05 |
| | Median | | $-0.79$ | 0.1 | 0.22 | 0.05 |
| | 95% | | $-0.5$ | 0.1 | 0.22 | 0.05 |
| (A4) | 5% | | $-1.26$ | 0.01 | 0.01 | 0.05 |
| | Median | | $-1.26$ | 0.05 | 0.05 | 0.05 |
| | 95% | | $-0.79$ | 0.1 | 0.1 | 0.1 |

Again, we also analyzed all twelve scenarios under the assumption that the observed SFS has converged to the expected value, to study the effect of model misspecification on parameter estimation. The results are shown in Table 5. The biases caused by incorrectly assuming *c*_0_ = *c_s_* are largest for the scenario that captures the youngest bottleneck (*t_S_* = 1/200). Here, not only the selection coefficients are strongly misestimated but also the time since the bottleneck (t^_*s*_) is largely underestimated. In all the other scenarios, at least the time since the bottleneck (t^_*s*_) is accurately estimated. The estimation accuracy of the other demographic parameters and selection coefficient increases with bottleneck age and the concomitant decreasing impact of the ancestral population size on the SFS. In summary, we note that assuming a too simplistic demographic model can lead to large errors in parameter estimation.

**Table 5.** Parameter estimation results based on the expected SFS when the ancestral population size *c*_0_ is incorrectly assumed to equal the current size *c_s_*, while the true model has *c*_0_ = 1/2 and *c_s_* = 1. SFS were computed for the following demographic scenarios and selection coefficients. In terms of the demography, a bottleneck model was assumed according to Figure 6 with parameters *c*_0_ = 1/2, *c_B_* = 1/10, *t_B_* = 1/10 and *t_s_* = 1/200, 1/20 or 1/2. The selection coefficients were chosen as *σ* = 0, −1/2 and −2. The parameter estimates were obtained according to the model assuming *c*_0_ = *c_s_* (the grid for the 4-parameter space being a subset of the grid for the 5-parameter space) and by assuming either selection or neutrality in each case.

| Selection coefficient | $\sigma = 0$ | $\sigma = -1/2$ | $\sigma = -2$ |
| --- | --- | --- | --- |
| Demographic model | $(\hat{\sigma}, \hat{c}_B, \hat{t}_B, \hat{t}_S)$<br>$(\hat{c}_B, \hat{t}_B, \hat{t}_S)$ | $(\hat{\sigma}, \hat{c}_B, \hat{t}_B, \hat{t}_S)$<br>$(\hat{c}_B, \hat{t}_B, \hat{t}_S)$ | $(\hat{\sigma}, \hat{c}_B, \hat{t}_B, \hat{t}_S)$<br>$(\hat{c}_B, \hat{t}_B, \hat{t}_S)$ |
| Bottleneck with $t_S = 1/200$ | $(-3.420, 0.023, 0.050, 0.001)$<br>$(0.224, 0.224, 0.011)$ | $(-0.171, 0.224, 0.224, 0.011)$<br>$(0.224, 0.224, 0.011)$ | $(-5.848, 0.023, 0.050, 0.001)$<br>$(0.023, 0.100, 0.001)$ |
| Bottleneck with $t_S = 1/20$ | $(-1.260, 0.050, 0.050, 0.050)$<br>$(0.100, 0.224, 0.050)$ | $(-2., 0.050, 0.050, 0.050)$<br>$(0.100, 0.224, 0.050)$ | $(-0.794, 0.100, 0.224, 0.050)$<br>$(0.100, 0.224, 0.050)$ |
| Bottleneck with $t_S = 1/2$ | $(-0.292, 0.224, 0.500, 0.500)$<br>$(0.224, 0.500, 0.500)$ | $(0, 0.050, 0.100, 0.500)$<br>$(0.050, 0.100, 0.500)$ | $(-2., 0.224, 0.500, 0.500)$<br>$(0.050, 0.224, 0.500)$ |

#### Testing a dataset of Drosophila melanogaster

Here, we apply our method to analyze a dataset which has been recently used to estimate the joint demographic history of several populations of *Drosophila melanogaster* (Duchen et al. 2013). The dataset consists of 12 sequences from a Zimbabwe population comprising 197 non-coding loci; and within each locus there are between 1 and 41 segregating sites (3234 polymorphic sites in total). We carry out our analysis based on the bottleneck model of the previous section assuming that the current and the ancestral population sizes are allowed to differ, assuming either neutrality or selection on the derived variant. Since purifying selection is assumed to be more prevalent than positive selection in intronic and intergenic regions of African *Drosophila*, we focus on a negative selection coefficient in our analysis.

We primarily use all segregating sites in our analysis. However, whereas the loci are scattered over the genome with at least tens of thousands of bases apart, the sites within each locus are tightly linked and hence are not independent. To study the effect of this discrepancy between the theoretical independence assumption underlying our method and the data, we also try using a subset of presumably independent sites by sampling one from each locus.

To begin with, a coarse maximum likelihood estimate of (*ĉ*_0_, *σ^*, *ĉ_B_*,*t*^_*B*_,*t^*_*S*_)=(1,0,0.05,0.1,0.1) was computed under the selective and the neutral bottleneck model on the parameter grid specified earlier. For each model, we investigated the accuracy of this parameter estimate via parametric bootstrap, using 200 bootstrap samples each consisting of 3234 polymorphic sites. Quantiles of the MLEs from the bootstrap samples are shown in Table 6, and, e.g., the confidence interval of the estimate of the ancestral population size (ĉ_0_) spans nearly the entire given parameter range.

**Table 6.** Parametric bootstrap results for the *Drosophila melanogaster* data The demographic history was estimated with and without selection for the demographic model illustrated in Figure 6 for the entire dataset of 3234 polymorphic sites. The estimation procedure is described in the main text. The estimated parameters given at the top of the table were used to generate 200 frequency spectra consisting of 3234 polymorphic sites each for the neutral and the selective model, respectively. We estimated the neutral demography for each neutral subsample and the demography with selection for each selective dataset, before the median, the 0.05 and the 0.95 quantiles were evaluated.

| | $\hat{\sigma}$ | $c_0$ | $\hat{c}_B$ | $\hat{t}_B$ | $\hat{t}_S$ |
| --- | --- | --- | --- | --- | --- |
| MLE | 0 | 1 | 0.05 | 0.1 | 0.1 |
| 5% | -0.794 | 0.5 | 0.002 | 0.002 | 0.1 |
|  |  | 0.5 | 0.023 | 0.023 | 0.1 |
| Median | -0.171 | 1 | 0.050 | 0.100 | 0.1 |
|  |  | 1 | 0.050 | 0.100 | 0.1 |
| 95% | 0 | 10 | 0.050 | 0.224 | 0.1 |
|  |  | 10 | 0.100 | 0.224 | 0.1 |

This suggests to improve the parameter estimates by successively refining the grid. The parameter range of each parameter was adjusted by choosing the respective two outermost parameter estimates from the set of the 100 likeliest parameter combinations of the coarse grid. We fixed the five possible ancestral population sizes *c*_0_ ∈ [0.5,10] (cf. Table 1) occurring in this set, and adopted a grid search for each of them, with *σ* ∈ [‒0.79,0], *c*_*B*_ ∈ [0.001,0.1], *t_B_*/*c_B_* ∈ [1,5] and *t_S_* ∈ [0.05,0.224]. Besides zero, 10 values were chosen for *σ*, 10 values for *c_b_*, and 30 values each for *t_B_*/*c_B_* and *t_S_*, so that a total of 99,000 grid points were applied for each c_0_. The ratio of two consecutive values in each parameter range is kept constant similarly to above. To focus on rescaled time *t_B_*/*c_B_* instead of *t_B_* relies on the observation that *t_B_* and *c_b_* correlate strongly and has the advantage that unlikely combinations of *t*_*B*_ and *c*_*b*_ can be omitted. More values were chosen for time parameters, since these are more sensitive than the population size parameters.

This procedure was repeated twice, upon which the maximum likelihood value did barely change. Each refined grid was based on the 100 likeliest parameter estimates, and the number of different possible ancestral population sizes was also successively raised to further refine the parameter *c*_0_. The maximum likelihood estimates for a range of parameters *c*_0_ and the associated likelihoods are given in Table 7. Selection is barely needed to explain the dataset and the estimated bottleneck population size (*ĉ_B_*) has reached the smallest possible value of 0.001 over the various grid searches. Choosing even distinctly smaller values for *ĉ_b_* would barely change the likelihood value anymore as long as the scaled bottleneck duration t^*_B_*/*_ĉ__B_* is kept constant. The time since the bottleneck (t^_*S*_) is robustly estimated over the various demographies that provide a similar likelihood (*L*), whereas the estimated bottleneck duration (*t^_B_*) correlates strongly with the ancestral population size (c_0_). Again, for each of the various ancestral population sizes (c_0_), the set of the 100 likeliest parameter combinations was used to obtain the parameter and likelihood ranges presented in Table 8. As one can see, most parameters were sufficiently pinpointed. In Figure 7, the SFS for the most likely neutral and selective parameter estimates, which can be found in the two penultimate lines of Table 7, are compared with the observed data.

**Table 7.** Estimated parameters and their likelihoods for the *Drosophila melanogaster* data for fixed values of *c*_0_ The demographic history was analyzed with and without selection for the demographic model illustrated in Figure 6 for the entire dataset of 3234 polymorphic sites. The maximum likelihood estimates and their likelihood values *L* were obtained for various predefined ancestral population sizes and based on a gradually refined grid as described in the main text.

| $c_0$ | $\hat{\sigma}$ | $\hat{c}_B$ | $\hat{t}_B$ | $\hat{t}_S$ | $L$ |
| --- | --- | --- | --- | --- | --- |
| 1.00 | 0 | 0.001 | 0.0015 | 0.1566 | −5963.888 |
| 1.67 | 0 | 0.001 | 0.0020 | 0.1614 | −5963.208 |
| 2.45 | 0 | 0.001 | 0.0024 | 0.1647 | −5963.027 |
| 3.59 | 0 | 0.001 | 0.0028 | 0.1614 | −5962.965 |
| 4.08 | 0 | 0.001 | 0.0029 | 0.1638 | −5962.924 |
| 5.99 | −0.007 | 0.001 | 0.0032 | 0.1655 | −5962.913 |
| 6.81 | −0.005 | 0.001 | 0.0034 | 0.1655 | −5962.902 |
| 7.74 | −0.002 | 0.001 | 0.0035 | 0.1655 | −5962.890 |
| 8.80 | 0 | 0.001 | 0.0037 | 0.1638 | −5962.884 |
| 10.0 | −0.007 | 0.001 | 0.0038 | 0.1647 | −5962.894 |

**Table 8.** Parameter ranges of the most likely estimates for the *Drosophila melanogaster* data for fixed values of *c*_0_ The demographic history was analyzed with and without selection for the demographic model illustrated in Figure 6 for the entire dataset of 3234 polymorphic sites. The set of the 100 likeliest parameter combinations was respectively estimated for various predefined ancestral population sizes and based on a gradually refined grid as described in the main text. From this set, the two outermost estimates were chosen for each single parameter and for the likelihood value *L* to obtain the outlined parameter ranges.

| $c_0$ | $\hat{\sigma}$ | $\hat{c}_B$ | $\hat{t}_B/\hat{c}_B$ | $\hat{t}_S$ | $L$ |
| --- | --- | --- | --- | --- | --- |
| 1.00 | $[-0.010, 0]$ | $[0.001, 0.0026]$ | $[1.49, 1.49]$ | $[0.1541, 0.1591]$ | $[-5963.926, -5963.888]$ |
| 1.67 | $[-0.007, 0]$ | $[0.001, 0.0021]$ | $[2.00, 2.00]$ | $[0.1598, 0.1630]$ | $[-5963.233, -5963.208]$ |
| 2.45 | $[-0.010, 0]$ | $[0.001, 0.0018]$ | $[2.37, 2.37]$ | $[0.1622, 0.1663]$ | $[-5963.050, -5963.027]$ |
| 3.59 | $[-0.015, 0]$ | $[0.001, 0.0026]$ | $[2.74, 2.77]$ | $[0.1598, 0.1680]$ | $[-5962.981, -5962.965]$ |
| 4.08 | $[-0.007, 0]$ | $[0.001, 0.0021]$ | $[2.89, 2.89]$ | $[0.1622, 0.1663]$ | $[-5962.949, -5962.924]$ |
| 5.99 | $[-0.015, 0]$ | $[0.001, 0.0021]$ | $[3.25, 3.25]$ | $[0.1622, 0.1688]$ | $[-5962.939, -5962.913]$ |
| 6.81 | $[-0.010, 0]$ | $[0.001, 0.0021]$ | $[3.38, 3.38]$ | $[0.1622, 0.1688]$ | $[-5962.926, -5962.902]$ |
| 7.74 | $[-0.007, 0]$ | $[0.001, 0.0018]$ | $[3.52, 3.52]$ | $[0.1622, 0.1680]$ | $[-5962.912, -5962.890]$ |
| 8.80 | $[-0.003, 0]$ | $[0.001, 0.0026]$ | $[3.66, 3.66]$ | $[0.1614, 0.1655]$ | $[-5962.914, -5962.884]$ |
| 10.0 | $[-0.022, 0]$ | $[0.001, 0.0026]$ | $[3.71, 3.75]$ | $[0.1614, 0.1688]$ | $[-5962.927, -5962.894]$ |

**Figure 7.**
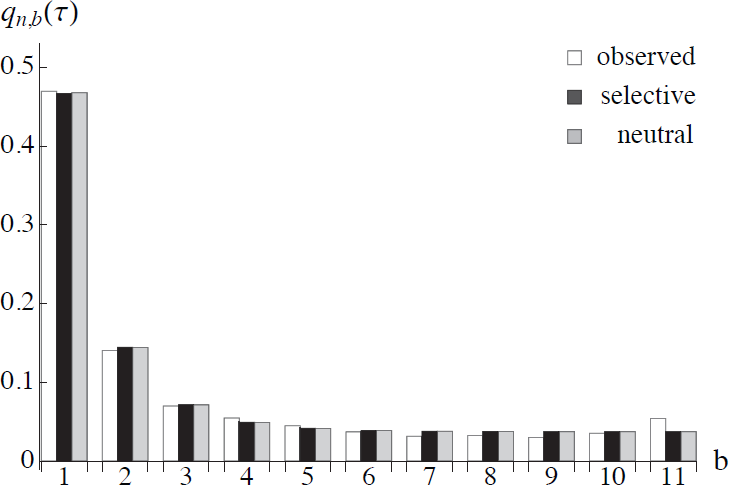
(a) SFS for the observed data and the most likely selective and neutral parameter estimates from left to right.

Comparing the SFS obtained using our parameter estimates and the ones given in Duchen et al. (2013), we obtain an improved goodness-of-fit to the observed SFS from the data. This is not surprising, since primarily statistics *summarizing* the SFS were used in their study. Furthermore, the authors allowed for different population sizes before and after the bottleneck as well but restricted the duration of the bottleneck to a somewhat arbitrary predefined value. The method in our work does not take the mutation rate explicitly into account, and thus cannot estimate the reference population size. Thus it would be too speculative to date the bottleneck in calendar time and to compare our outcome to the estimate of Duchen et al.

To investigate the effect of linkage within each of the 197 sequence fragments in the original dataset, we sampled one site per fragment to obtain a dataset consisting of 197 polymorphic sites. We repeated this procedure 200 times and maximized the log-likelihood for each sample similarly as above. Compared to the analysis of the full dataset, the SFS computed from the median parameter estimate shows a poorer fit to the data. This is likely due to the strong stochasticity in the bootstrap resampling procedure, since the individual parameter estimates for each sample do provide a good fit despite the small number of sites considered.

It might be tempting to assume that the excess of high-frequency derived variants in the observed data might be a result of weak positive selection. Therefore, we conducted an equivalent analysis as above, starting from the same grid with inverted signs for the selection coefficients. However, we did not obtain estimates being plausible from a biological point of view, since the estimation procedure favours selection coefficents in the upper range of the chosen interval [0,10]. When, as in this example, an excess of low- and high-frequency derived variants is simultaneously observed in comparison to a standard neutral model, unrealistically large estimates for *σ* are needed to explain the data. Positive selection on its own (and of some appreciable strength) causes a decline of low-frequency derived variants and an excess of high-frequency derived alleles, whereas an expansion (as embedded in the bottleneck model) acts vice versa. Therefore, both forces have to severely counteract each other so that the requirements of both ends of the SFS can be met.

### A model of human exponential population growth

We now demonstrate the utility of our method to investigate population size histories containing epochs of exponential growth in combination with selection. To this end, we adopt the following demographic history of a sample of African human exomes that has been estimated by Tennessen et al. (2012) as a modification of a model by Gravel et al. (2011). The population had an ancestral size of 7310 individuals until 5920 generations ago (assuming a generation time of 25 years), when it increased instantaneously in size to 14,474 individuals. After this increase, the population remained constant in size until 205 generations ago, when it started to grow exponentially until reaching 424,000 individuals at present. The relative population size function for this model can be described by

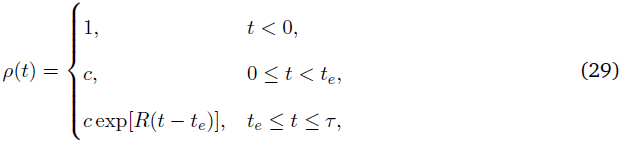

where *c* is the ratio of population sizes after and before the instantaneous expansion, which can be dated arbitrarily, so we set the time of this expansion to zero. *R* is the scaled exponential growth rate, *t_e_* is the time at which the expansion started, and *τ* is the time of sampling (the present). Times are given in units of 2*N*_ref_, where the reference population size *N*_ref_ is the initial size before time zero (the ancestral size). Since the theoretical framework presented above assumes a history of piecewise constant population sizes, the phase of exponential growth in this model has to be adequately discretized to obtain a suitable piecewise approximation. The following piecewise function can be chosen to approximate the exponential growth phase via a geometric growth function:

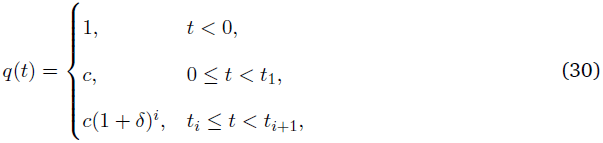

with times *t_i_* = *t_e_* + log [(1 + *δ*)^*i*−1^(2 + δ)/2] /*R*, *i* = 1,…, *i_τ_*. Here, the number of population size changes during the phase of exponential growth is given by

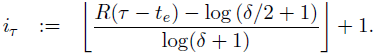

Varying the growth rate *δ* determines the number of discretization intervals used.

The SFS (28) of the discretized version is obtained straightforwardly from (26) and (27). For the demographic parameters given above, we computed the SFS for various sample sizes up to 200 and we used *δ* = 1/4, which was chosen large enough to provide reasonable fast computation times but sufficiently small to provide a good approximation of the exponential growth model. In the neutral case, the goodness of the approximation can be verified via the explicit solution of the SFS (Živković and Stephan 2011), which can be applied to the continuous and the discretized model. As shown in Figure 8a, where a sample size of *n* = 200 is chosen, the spectra of both continuous and piecewise-constant models agree very well with each other; the percentage error is 0.57% based on the *l*^2^-norm, while the Kullback-Leibler divergence is about 1.76 × 10^−7^.

**Figure 8.**
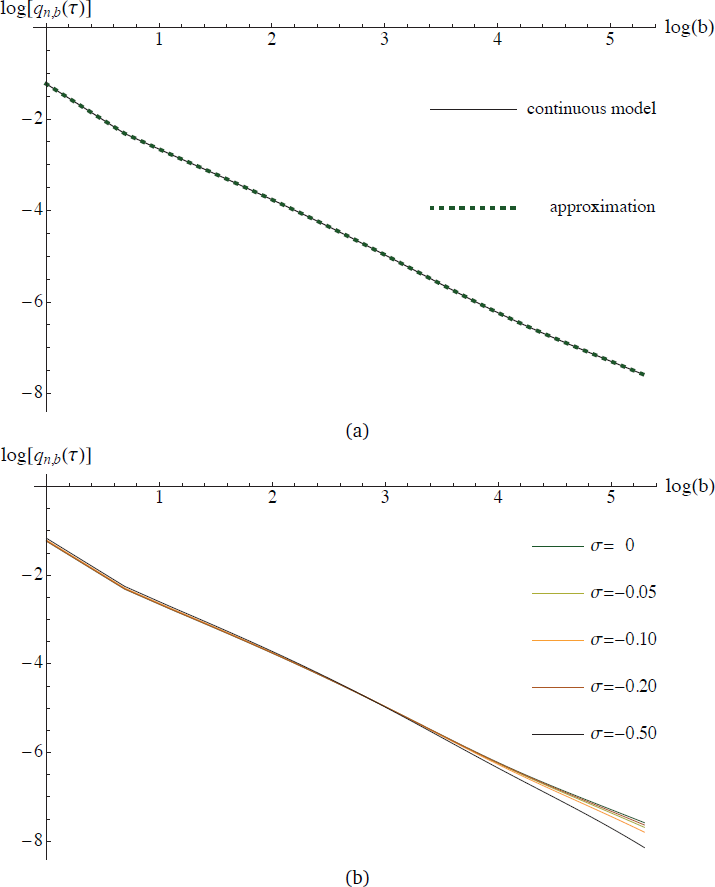
(a) Log-log plots for the SFS of the continuous and the discretized version of the estimated human African demography and neutral evolution. (b) Log-log plots for the SFS of the discretized version under various selection coefficients. The selection coefficients in the legend are ordered from top to bottom according to the function values of the high-frequency derived alleles. The sample size is given by *n* = 200 in both subfigures.

Using our method, selection can then be incorporated into the piecewise-constant population size model. The effect of various negative selection coefficients (scaled with respect to the ancestral population size) is illustrated again for sample size *n* = 200 in Figure 8b, and the same trend can be observed for smaller sample sizes as well. It is probably not surprising that the resolution in distinguishing the selective and the neutral model rises with *σ*. More interestingly, differences between the neutral and the selective models are apparently more pronounced among derived alleles in intermediate- to high frequency. Therefore, for large datasets where intermediate to high-frequency derived alleles are present in sufficient numbers, one may focus more strongly on these allelic classes than on low-frequency derived ones for the statistical analysis of purifying selection.

## Discussion

Already in the early days of population genomics, several studies in various species (e.g.,Glinka et al. 2003, Williamson et al. 2005) have revealed that both natural selection and demographic forces have shaped the patterns of polymorphism in modern samples of DNA sequences. However, most inference methods relied on computer simulations and the usage of statistics that have been designed to detect deviations from neutrality assuming a constant population size (e.g., Glinka et al. 2003). More elaborate approaches utilized the transition density function that describes allele frequency changes over time, where most methods solved the underlying diffusion equation numerically employing discretization schemes (e.g., Williamson et al. 2005, Zhao et al. 2013). Besides several issues that may arise in such purely numerical frameworks, only the simplest models of a single population size change have been considered in these studies. Recently, Song and Steinrücken (2012) developed a more analytical approach that provides the spectral representation of the transition density for a model that includes general diploid selection and recurrent mutations under a constant population size.

In this article, we extended their solution for the case of genic selection to an arbitrary number of instantaneous changes in population size. First, we obtained a rescaled version of the spectral representation of the transition density function for a single time period during which the population size differs with respect to a reference size. Combining the transition densities for single time periods over arbitrarily many time points of instantaneous population size changes yields the transition density function for such a multi-epoch model with genic selection.

The transition density function has been employed to obtain the SFS. However, explicit knowledge of the transition density function is not required for the computation of the SFS. We revisited and simplified a method by Evans et al. (2007) who expressed the allele frequencies in terms of their moments. Their method requires that a system of ordinary differential equations is solved numerically. They employed a finite difference scheme to tackle a model of strong exponential growth with genic selection. We simplified this approach starting from a model of a constant population size, to which instantaneous population size changes were recursively added. The numerically obtained result at the end of a certain epoch is used as the initial condition of the subsequent time phase. This simple procedure allows us to consider numerous population size changes and offers fast computations with little loss in accuracy. We have shown that even a model of strong population growth can be well approximated.

The transition density function with variable population size can be incorporated into a hidden Markov model framework (cf. Steinrücken et al. 2014 for a constant population size) to analyze time series genetic data. However, in this article we focused on several biological questions that can be investigated using the SFS, and treat the time series application in a separate paper. We first addressed the joint estimation of bottleneck and selection parameters from polymorphism data within a maximum likelihood framework. This approach can be applied to *simultaneously* infer selection coefficients and the parameters of a model of instantaneous population size changes. The importance of methods that allow to estimate selective and demographic parameters jointly becomes particularly apparent in large populations for which the scaled selection coefficient, *σ*, can take considerable values across large regions of the genome, so that demography and selection cannot be estimated independently. Although selection is known to act either positively or negatively and with different strengths across the genome, a constant selection coefficient has been applied in our approach. A constant selection coefficient can either be interpreted as a genome-wide average, or, more realistically, as the selection strength of a certain functional class, among which the coefficients should not vary greatly. This argument particularly applies to the *Drosophila* example for which intronic and intergenic loci were sequenced and used for the parameter estimation.

For the first part of *Applications*, we generated data for the estimation procedure by sampling a large number of sites from the SFS of a bottleneck model varying the strength of selection. We assumed the same and also a slightly less complex model with five and four free parameters, respectively, for the parameter estimation. We demonstrated that our method can accurately estimate the parameters in the majority of the bottleneck scenarios, but less so, when the simpler model is assumed. The time since the bottleneck was retrieved in most of the cases even when assuming the simpler model. It is interesting to note that even when the datasets simulated with selection are analyzed assuming neutrality, the time since the bottleneck was quite robustly inferred except for the briefest one being estimated under the simpler 4-parameter model. This result is quite promising with respect to the many published demographic estimates that have been obtained assuming neutrality, because the time since the last demographic change might not be subject to major changes. However, this result has to be investigated further in more realistic models that also include phases of exponential growth, which can be studied based on our results as well. Our results encourage the application of rather complex than too simple demographic models anyway.

In the African *Drosophila* sample, no or barely any negative selection was inferred, which might simply be a result of well chosen neutral markers that barely experienced selection. Furthermore, it turned out to be difficult to pinpoint in particular the ancestral population size and the duration of the bottleneck, whereas the time since the bottleneck was robustly estimated. From a theoretical point of view, Bhaskar and Song (2014) have recently obtained sufficient conditions for the identifiability of piecewise-defined demographic models under neutrality using the expected frequency spectrum; the identifiability of demographic models combined with selection is an interesting future research question. However, one has to keep in mind that the estimates were obtained from partly linked loci of a small sample of chromosomes and that taking a subset of independent loci to meet the theoretical assumptions result in relatively small datasets showing large variance in the estimates.

We finally analyzed an example of exponential human population growth (Tennessen et al. 2012) to see the effect of purifying selection in the context of this model. As illustrated in Figure 8b for a sample of size 200 and various selection coefficients, intermediate- and high-frequency derived variants are more affected by exponential growth and negative selection than the low-frequency derived ones. A plausible reason is that both exponential growth and negative selection enforce an increase of low-frequency derived variants until these classes are saturated and their impact can rather be observed in the complimentary high-frequency allelic classes. In general, this example illustrates nicely that even more elaborated models that include various phases of exponential growth and population declines can be computationally efficiently treated via an appropriate discretization of phases of continuous population size change, using the methods presented in this paper.

## Acknowledgements

We thank the generous support of the Simons Institute for the Theory of Computing, where much of this work was carried out while we were participating in the 2014 program on “Evolutionary Biology and the Theory of Computing.” DZ thanks Anand Bhaskar, Steven N. Evans and Andreas Wollstein for helpful discussions. YSS thanks the Miller Institute for providing a Research Professorship while this paper was completed. This research is supported in part by DFG grant STE 325/14 from the Priority Program 1590 (DZ, WS), the Volkswagen Foundation grant I/84232 (DZ), an NIH grant R01-GM094402 (MS, YSS), and a Packard Fellowship for Science and Engineering (YSS).

## Appendix. Derivation of (12)

Here, we derive the expression shown in (12). Using (2), (5), and (7), note that

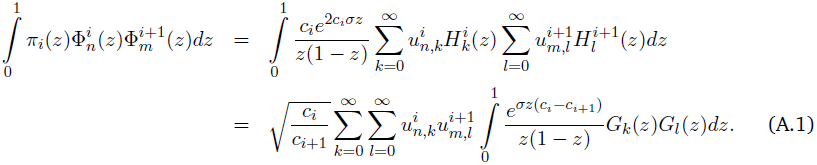

Without loss of generality, assume *c*_*i*_ ≠ *c*_*i*+1_. (If *c*_*i*_ = *c*_*i*+1_, the integral in (A.1) is trivial to evaluate using orthogonality.) Since *z*^−1^(1 – *z*)^−1^*G*_*k*_(*z*)*G*_*l*_(*z*) is a polynomial of order *k* + *l* + 2, its *j*th derivative vanishes for j ≥ *k* + *l* + *3*. Using integration by parts recursively *k* + *l* + 2 times, we obtain

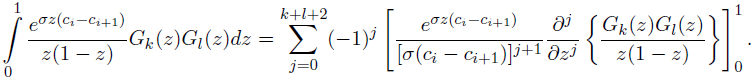

Note that the summand for *j* = 0 in the previous equation is equal to zero and will be omitted in the remainder. Since *G*_*k*_(1 – *z*) = (−1)^*k*^*G*_*k*_(*z*), we have

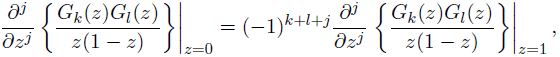

so that

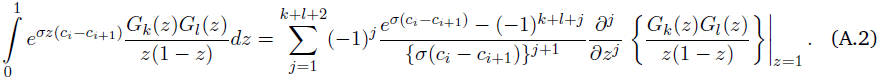

The modified Gegenbauer polynomials are defined as

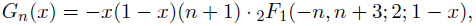

where 
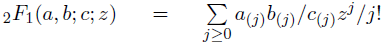 is the Gauss hypergeometric function, d(0) = 1, and *d*_(*j*)_ = *d*(*d* + 1) … (*d* + *j* – 1), j ≥ 1. Applying this definition, we obtain

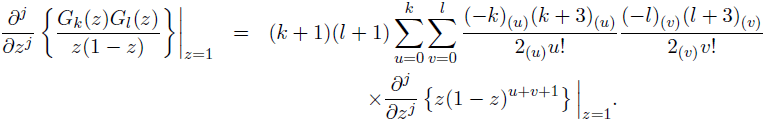

Note that the sums are finite, since (−*a*)(*b*) = 0 for integers *a* < *b*. It is simple to show that

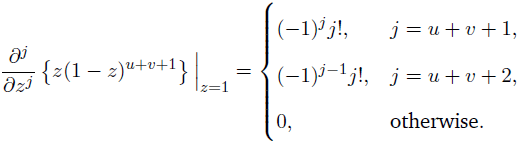

By applying this result we obtain, after some algebra,

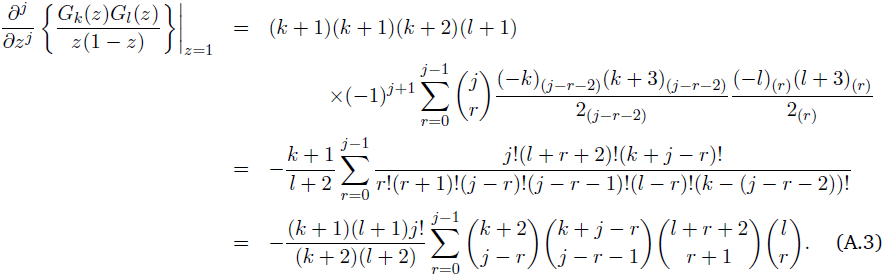

Finally, combining (A.3), (A.2), and (A.1) yields the desired result.

